# Sex Differences and Role of Lysyl Oxidase Like 2 (LOXL2) in Angiotensin II-Induced Hypertension in Mice

**DOI:** 10.1101/2023.12.13.571541

**Authors:** Huilei Wang, Marta Martinez Yus, Travis Brady, Rira Choi, Kavitha Nandakumar, Logan Smith, Rosie Jang, Bulouere Princess Wodu, Jose Diego Almodiel, Laila Stoddart, Deok-Ho Kim, Jochen Steppan, Lakshmi Santhanam

## Abstract

**Background:** Hypertension, a disease with known sexual dimorphism, accelerates aging associated arterial stiffening, in part due to the activation of matrix remodeling caused by increased biomechanical load. In this study, we tested the effect of biological sex and the role of the matrix remodeling enzyme lysyl oxidase like 2 (LOXL2) in hypertension induced arterial stiffening.

**Methods:** Angiotensin II (Ang II) was delivered using osmotic pumps in *Loxl2^+/-^* and WT male and female mice. Blood pressure and pulse wave velocity (PWV) were measured noninvasively to assess hypertension and aortic stiffness. Wire myography and uniaxial tensile testing were used to test aortic vasoreactivity and mechanical properties. Aortic wall composition was examined by histology and Western blotting. The effect of biomechanical strain on LOXL2 expression and secretion by vascular smooth muscle (VSMC) and endothelial cells (EC) was evaluated by uniaxial cyclic stretching of cultured cells. The role of LOXL2’s catalytic function on VSMC alignment in response to mechanical loading was determined with LOXL2 inhibition and knockout.

**Results:** Ang II infusion induced hypertension in WT and *Loxl2^+/-^* mice of both sexes and increased PWV in WT males but not in *Loxl2^+/-^* males, WT females, or *Loxl2^+/-^* females. LOXL2 depletion protected males from Ang II mediated potentiation of vasoconstriction but worsened in females and improved aortic mechanical properties in both sexes. Histological analysis showed increased aortic wall thickness in hypertensive WT males but not females and increased intralamellar distance in both sexes, that was ameliorated in Loxl2+/- mice. Western blotting revealed increased collagen I, decreased collagen IV, and increased LOXL2 accumulation and processing in hypertensive mice. Hypertensive cyclic strain contributed to LOXL2 upregulation in the cell-derived matrix in VSMCs but not ECs. LOXL2’s catalytic function facilitated VSMC alignment in response to biomechanical strain.

**Conclusions:** In males, arterial stiffening in hypertension is driven both by VSMC response and matrix remodeling. Females exhibit a delayed onset of Ang II-induced hypertension with minimal ECM remodeling but with VSMC dysfunction. LOXL2 depletion ameliorates arterial stiffening and preserves functional contractility and aortic structure in male hypertensive mice. LOXL2 depletion improves aortic mechanics but worsens aortic contractility in hypertensive females. VSMCs are the primary source of LOXL2 in the aorta and hypertension increases LOXL2 processing and shifts to collagen I accumulation. Overall, LOXL2 depletion offers protection in young hypertensive males and females.

## Introduction

Hypertension is one of the major risk factors for cardiovascular diseases, including coronary artery disease, cardiac hypertrophy, atrial fibrillation, stroke, heart failure, and renal failure^1–6^. The global prevalence of hypertension has been increasing in the past four decades, to an estimated 31.1% in the global adult population^7^. Though awareness and treatment of hypertension has improved, only 43.4% of US patients with documented hypertension are able to successfully manage high blood pressure^8^. One of the consequences of uncontrolled hypertension is the acceleration of arterial stiffening noted with natural aging^9^. Because aortic stiffening is both a cause and a consequence of hypertension^10^, an insidious loop of stiffening occurs, and this exacerbates cardiovascular deterioration. Therefore, it is of high clinical interest to target arterial stiffening in the context of hypertension.

Population level data demonstrate remarkable sexual dimorphisms in both hypertension and age-related arterial stiffening. Young women have similar (32% in women vs 34% in men) ^11^ or even lower (13% in women vs 31.2% in men)^12^ rates of diagnosis, treatment, and control. However, morbidity and mortality due to cardiovascular events remains higher in young to middle-aged men. Estrogen has been widely studied for its beneficial effects in young women. The roles of estrogen in maintaining NO bioavailability^13, 14^ and vascular function^15^ are well documented. However, estrogen supplementation in post-menopausal females (human and preclinical models) has shown mixed results^16–18^. This points to the involvement of additional mechanisms in guiding the sex differences in hypertension.

Aortic stiffening arises with aging, and hypertension is known to accelerate arterial stiffening during aging. Remodeling of the vascular medial extracellular matrix (ECM) by involving increased deposition and crosslinking of collagens by the resident vascular smooth muscle cells (VSMC) is one of the key components of arterial stiffening induced by hypertension^19–22^. While this is initially an adaptive and necessary response, particularly in the resistance vessels to normalize wall tension in the face of higher blood pressure, the process is eventually maladaptive, particularly when remodeling of the compliance vessels (such as the aorta) is noted. Whether young females exhibit lower ECM remodeling in response to hypertension when compared with males is not known.

In addition to the synthesis and secretion of structural proteins, the concerted activation of matrix remodeling enzymes is necessary for structural remodeling to occur. The lysyl oxidase (LOX) family of amine oxidase enzymes are central to matrix stability, homeostasis, and remodeling. Of these, LOX and LOX-like 2 (LOXL2) have been shown to be critically important to cardiovascular function and play a role in disease progression^23–25^. Prototypic LOX function is critical for structural stability of the aorta even in the absence of hypertension, as evidenced by aortic dilation and aneurysms with LOX polymorphisms associated with loss of its function.^26, 27^ In pre-clinical models of hypertension, LOX inhibition (e.g. with the small molecule inhibitor β- aminopropionitrile (BAPN)), leads to an increased incidence of aortic aneurysms and increased rupture^28, 29^. Indeed, this has led to Ang II+BAPN being a well-accepted experimental model of abdominal aortic aneurysm (AAA) in mice and rats.^30–33^ Thus, LOX itself is not a suitable target to pursue in the context of vascular remodeling in aging or hypertension. On the other hand, LOXL2, a member of the LOX family, has emerged as an important and promising target in cardiovascular disease. LOXL2 has been shown to play a role in the development of various cardiovascular diseases including cardiac interstitial fibrosis, atrial fibrillation, and cerebral aneurysm.^23, 24, 34–36^ Serum LOXL2 was shown to be elevated in heart failure patients and increased LOXL2 expression was noted in human hearts following ischemic or nonischemic dilated cardiomyopathy.^24^ Prior studies have identified LOXL2 as a potential therapeutic target in aging associated vascular stiffening^37^ and shown higher levels of LOXL2 activity in aortic extracellular matrix in aged mice.^37, 38^ Importantly, LOXL2 depletion in mice delayed aging associated aortic stiffening without any deleterious effects (i.e., no increase in aortic tortuosity, dilation, or aneurysm formation).^37^

In hypertension, augmented biomechanical strain experienced by the vascular cells due to increase in blood pressure is a major contributor to activation of ECM remodeling.^39, 40^ Mechanical forces have been shown to cause in changes to VSMC function (e.g. contractility of VSMCs) and behavior (e.g. proliferation), and structural changes (ECM remodeling and stiffening). While prior studies have shown that cyclic strain mimicking hypertension increases matrix production by VSMCs^39^, the effect of biomechanical strain on LOXL2 regulation and the specific cellular source in the vessel wall (VSMCs or ECs) are not known.

Additional elements of LOX/LOXL2 biochemistry can also contribute to hypertension induced vascular remodeling. First, LOX is secreted as a pro-protein (∼50 kDa) and requires processing by BMPs to release the active form (∼25 kDa)^41^. On the other hand, LOXL2 is active in its full-length form (∼100 kDa), and processing shifts its substrate preference from collagen IV for the full-length form to collagen I for the processed form (∼65 kDa)^42^. Thus, both synthesis/secretion and processing are putative mechanisms by which LOX and LOXL2 could be regulated in the hypertensive vasculature. Whether LOX and LOXL2 are proteolytic processing contributes to shifts in collagen subtype content in the hypertensive aortic ECM is not known. Second, while ECM remodeling by LOX/LOXL2 and the resultant change in collagen subtype content can contribute to shifts in cell alignment/behavior arising from mechanosensing, LOXL2 also contains four scavenger receptor cysteine rich (SRCR) domains that suggest catalytically independent functions arising through protein motif recognition, particularly in mediating cell-ECM interactions related to cell alignment. The putative catalytically-independent roles of LOXL2 is therefore important to discern.

There is a clear effect of sex in aging-associated arterial stiffening. Specifically, women are relatively protected from arterial stiffening during menstruating years but exhibit a marked acceleration in the post-menopausal years.^43, 44^ In young individuals, potential differences in the temporal evolution of arterial stiffening with hypertension, similar to aging (i.e., whether hypertensive females exhibit a dampened stiffening response when compared with hypertensive males) is not known. Further, sex differences in the cellular and molecular mechanisms that mediate structural changes in the aorta in response to hypertension remain incompletely understood.

Therefore, given the sexual dimorphism of aortic stiffening during aging and hypertension, the role of LOXL2 in aortic stiffening during aging, and the regulation of matrix remodeling by biomechanical strain, we tested the hypotheses that: 1) young females are relatively protected from hypertension induced aortic stiffening when compared to young male counterparts, 2) LOXL2 activation by biomechanical strain contributes to ECM remodeling in hypertension and 3) LOXL2 depletion is protective against hypertension induced aortic stiffening. Within this context, we also evaluated if ECs and/or VSMCs are the source of LOXL2, whether LOX/LOXL2 proteolytic processing is a mechanism by which aortic matrix remodeling is regulated in hypertension, and whether LOXL2 contributes to cell-alignment independent of its catalytic function.

## Methods

### Animal model

LOXL2 gene was disrupted by inserting a long terminal repeat (LTR)-trapping cassette in exon 1 as previously described^37^. *Loxl2^+/−^* mice (generated on a C57Bl/6J background) were bred in-house in pairs and genotyped by PCR of tail biopsy. Male and female *Loxl2^+/−^*were used in this study as *Loxl2*^-/-^ is perinatal lethal.^45^ Littermate *Loxl2^+/+^* (Wild type; WT) were used controls. Additional age-matched C57BL/6J mice were purchased from the Jackson Laboratory to supplement the male and female WT groups. All procedures involving animals were approved by the Institutional Animal Care and Use Committee of the Johns Hopkins University. Mice used in the study were housed in groups (5 mice per cage; grouped by sex, litter, and study group (i.e., ±Ang II)) in the Johns Hopkins University School of Medicine animal care facility, fed and watered ad libitum in a temperature-controlled room with a 12:12-hour light/dark cycle. For longitudinal studies, mice were handled by the same individual and the longitudinal measures (described next) were acquired by the same individual during the same part of the day (8-10 am).

### Blood pressure (BP) measurement

BP was measured non-invasively in conscious mice using the CODA noninvasive tail-cuff BP system (Kent Scientific Corporation). Mice were first gradually acclimated to being restrained in the mouse holders over 10 days. Mice were maintained at 37 °C during measurement. BP was measured at baseline (immediately preceding Ang II pump placement), and then weekly after initiation of Ang II infusion. At least 5 measurements were recorded for each mouse for each time point, and then averaged.

### Pulse-wave velocity (PWV) measurement

PWV, the gold-standard index of in vivo arterial stiffness, was measured noninvasively using high-frequency Doppler (Indus Instruments) as previously described^46^. Briefly, mice were placed supine on a heated (37 °C) pad under anesthesia with 1.5% isoflurane. Pulsed Doppler signals and ECG were simultaneously captured at a thoracic and an abdominal aortic site. PWV (m/s) was calculated as the distance between these two locations, divided by the pulse transit time calculated using the foot-to-foot method.

### Hypertension model

Hypertension was induced by Angiotensin II infusion (Ang II; 1000 ng/kg/min) via osmotic mini pumps (Durect Corp) in male and female WT and *Loxl2^+/-^* mice (**Fig 1A**, n = 11-12 per cohort) as follows: After baseline BP and PWV measurement, 12-14 weeks old WT and *Loxl2^+/-^* mice were randomized into hypertension and control groups. For pump implantation, mice were anesthetized with 1.5% isoflurane and underwent subcutaneous implantation of osmotic minipumps filled with Ang II solution in PBS (Sigma-Aldrich). For analgesia, mice received acetaminophen-treated drinking water (1.6 – 3.2 mg/ml) for 24 hours prior surgical procedure and for 2 days after the procedure. Controls underwent sham surgery. BP and PWV were measured weekly after surgery for 3 weeks. At the end of the study period (3 weeks of Ang II infusion), body weight was measured, and the aorta and heart were dissected out. The heart was weighed, and heart weight/body weight was calculated as an index of myocardial hypertrophy. The aorta was cleaned free of connective tissue and perivascular adipose tissue, and then cut into 2-mm rings, and used immediately for tensile testing (2 intact and 2 decellularized rings) and wire myography (2 rings), fixed in 4% formaldehyde for histology (1 ring), or flash-frozen for Western blotting (1 ring).

**Figure 1.**
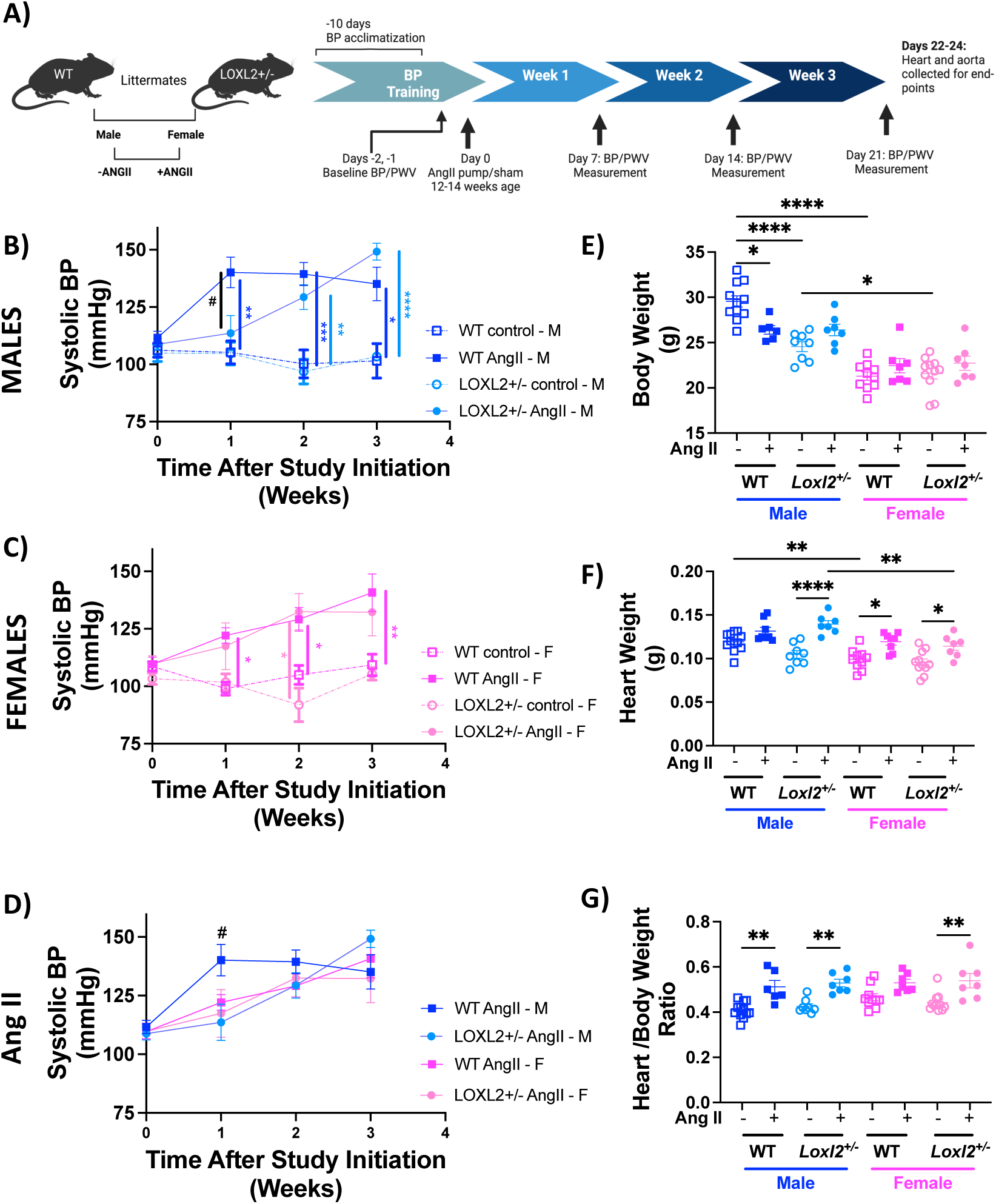
Angiotensin II (Ang II) infusion induced hypertension in males and females. **(A)** Schematic showing experimental timeline and endpoints. **(B)** Time course of systolic blood pressure (BP) of male (M) WT and *Loxl2^+/-^* mice with and without Ang II infusion (n = 8 for WT control M, n = 8 for WT + Ang II M, n = 8 for *Loxl2^+/-^* control M, and n = 7 for *Loxl2^+/-^* +Ang II M. At 1 wk: \*\**p* < 0.01 for WT Ang II M vs. WT control M and # p<0.05 for WT + Ang II M vs. *Loxl2^+/-^* +Ang II M; at 2 wks: \*\*\**p* < 0.001 for WT Ang II M vs. WT control M, \*\**p* < 0.01 for *Loxl2^+/-^* Ang II M vs. *Loxl2^+/-^* control M; at 3 wks: \**p* < 0.05 for WT Ang II M vs. WT control M, \*\*\*\**p* < 0.0001 for *Loxl2^+/-^* Ang II M vs. *Loxl2^+/-^* control M, by repeated measures 2-way ANOVA with Tukey *post hoc* analysis). **(C)** Time course of systolic BP of female (F) WT and *Loxl2^+/-^* mice with and without Ang II infusion (n = 10 WT control F, n = 8 WT+Ang II F, n=8 *Loxl2^+/-^* control F, n = 8 and *Loxl2^+/-^* +Ang II F. At 1 wk: \**p* < 0.05 for WT Ang II F vs. WT control F; at 2 wks: \**p* < 0.05 for WT+Ang II F vs. WT control F, \**p* < 0.05 for *Loxl2^+/-^* + Ang II F vs. *Loxl2^+/-^* control F; at 3 wks: \*\**p* < 0.01 for WT Ang II F vs. WT control F, by repeated measures 2-way ANOVA with Tukey *post hoc* analysis. **(D)** Time course of systolic BP of male (M) and female (F) mice with Ang II infusion (n = 8 for WT + Ang II M; n = 7 for *Loxl2^+/-^*+Ang II M; n = 8 WT+Ang II, F; n = 8 and *Loxl2^+/-^* +Ang II F; at 1 week, #p<0.05 WT+Ang II M vs. *Loxl2^+/-^* + Ang II M and WT+Ang II M vs. *Loxl2^+/-^* + Ang II F by 2-Way ANOVA with Tukey *post-hoc* analysis**) (E)** body weight of male and female WT and *Loxl2^+/-^* mice at the end of the experimental period (n= 12 for WT control M, n = 6 for WT+Ang II M, n = 8 for *Loxl2^+/-^* control M and n = 7 for *Loxl2^+/-^* + Ang II M, n= 10 for WT control F, n = 7 for WT+Ang II F, n = 11 for *Loxl2^+/-^* control F and n = 7 for *Loxl2^+/-^*+ Ang II F; * p< 0.01, ****p<0.0001 by 2-way ANOVA with Bonferroni *post hoc* analysis). **(F)** heart weight of male and female WT and *Loxl2^+/-^* mice at the end of the experimental period (n= 12 for WT control M, n = 6 for WT+Ang II M, n = 8 for *Loxl2^+/-^* control M and n = 7 for *Loxl2^+/-^* + Ang II M, n= 10 for WT control F, n = 7 for WT+Ang II F, n = 11 for *Loxl2^+/-^* control F and n = 7 for *Loxl2^+/-^* + Ang II F; * p< 0.01, ** p<0.01, ****p<0.0001 by 2-way ANOVA with Tukey’s *post hoc* analysis). **(G)** heart weight/ body weight ratios as an index of cardiac hypertrophy at the end of 3 weeks of the study period (n= 12 for WT control M, n = 6 for WT+Ang II M, n = 8 for *Loxl2^+/-^* control M and n = 7 for *Loxl2^+/-^*+ Ang II M, n= 10 for WT control F, n = 7 for WT+Ang II F, n = 11 for *Loxl2^+/-^* control F and n = 7 for *Loxl2^+/-^* + Ang II F; *p<0.05, ** p< 0.01, ****p<0.0001 by 2-way ANOVA with Tukey’s *post hoc* analysis).

### Wire myography

Vasoreactivity of the aortic rings was examined as previously described^37, 47^. Briefly, freshly dissected aortic rings (2 from each mouse) were mounted in the chamber of a wire myograph (DMT) in oxygenated Krebs-Ringer bicarbonate solution (in mM: 118.3 NaCl, 4.7 KCl, 1.6 CaCl2, 1.2 KH2PO4, 25.0 NaHCO3, 1.2 MgSO4, and 11.1 dextrose) at 37 °C and pre-stretched to 600 mg-force. Maximal contractility elicited by potassium chloride (KCl) was measured. After washing, vasocontractile response to increasing concentrations of phenylephrine (PE; 10^−9^–10^−5^Lmol/L) was measured. Next, endothelium-dependent (acetylcholine; ACh; 10^−9^–10^−5^Lmol/L) and endothelium-independent (sodium nitroprusside; SNP, 10^−9^–10^−5^Lmol/L) vasorelaxation of vessels preconstricted with PE (10^-6^Lmol/L) was determined. All reagents used in wire myography were purchased from Sigma Aldrich.

### Tensile Testing

The mechanical properties of the intact and decellularized aortic rings were determined by uniaxial tensile testing as previously described^48–51^. Two intact and two decellularized aortic rings from each mouse were tested. Samples were decellularized following published protocols^48^ with minor modifications. Briefly, aortic rings were placed in a decellularization solution of 1% SDS and 3.3% NH_4_OH in H_2_O, and gently rocked for 1 h at room temperature on an end-to-end shaker. Samples were then washed with 3.3% NH_4_OH in H_2_O (30 min, gentle rocking), followed by 3 washes (15 min each) in phosphate buffered saline (PBS, pH 7.4; ThermoFisher; containing antibiotic-antimycotic (1x, ThermoFisher)). Removal of cells was verified by the absence of DNA (Pico Green assay kit; Invitrogen), and the absence of GAPDH in the aorta homogenate by Western blotting.

For tensile testing, vessel lumen diameter (*Di*), wall thickness (*t*), and sample length (*L*) were first determined by microscopy for each sample. Samples were mounted on the pins of an electromechanical puller (DMT560; Danish Myo Technology A/S). After calibration and alignment, the pins were moved apart using an electromotor and displacement and force (*F*) were recorded continuously until sample failure. Engineering stress (*S*) was calculated by normalizing force (*F*) recorded with stress-free ring cross-section area (*S*□*=*□*F/2t*□*×*□*L*). Engineering strain (λ) was calculated as the displacement divided by *Di*. The stressLstrain relationship was represented by the equation S = α exp(βλ) where α and β are constants. α and β were determined by nonlinear regression for each sample and used to generate stress-strain curves.

### Cell Culture

Human aortic smooth muscle cells (HASMCs, ThermoFisher) were cultured in smooth muscle cell media (ScienCell) containing 2% fetal bovine serum, SMC-growth supplement, and antibiotic-antimycotic. LOXL2-depleted HASMC (T1) cells were generated by targeting the *Loxl2* gene by CRISPR-Cas9 gene editing as previously described^38^. Human Aortic Endothelial Cells (HAECs) were cultured using complete endothelial cell media (ECM; ScienCell) with 5% FBS, endothelial cell growth supplement antibiotic-antimycotic. All cells were cultured in humidified incubators with 5% CO_2_ at 37°C. To obtain LOXL2 enriched media, LOXL2 was overexpressed by adenoviral transduction as previously described. Prior to use, cells were authenticated by immunofluorescent staining of α-SMA and SM22α for HASMCs and CD31 (PECAM) and VEGF-R2 for HAECs. For both cell types, 3 distinct lot numbers (representing 3 distinct donors; 50-55 year old Caucasian males with no reported co-morbidities (hypertension, type 2 diabetes, obesity)) were used.

### Uniaxial cell stretching

To examine the effect of biomechanical strain on LOXL2 expression, HASMCs and HAECs were subjected to uniaxial cyclic mechanical stimulation at normotensive (10%) and hypertensive (20%) strain^52, 53^ using the Strex Cell Stretching System as follows: Cells were seeded on fibronectin coated PDMS chambers (150,000 cells/chamber for HASMCs, and 680,000 cells/chamber for HAECs). Each chamber has 4cm^2^ culture area. Cells were allowed to adhere and spread overnight and then serum starved for 24 h, following which fresh media was provided to the cells just before initiating uniaxial cyclic loading (10% or 20% at 1 Hz frequency). Static controls were maintained in the same incubator. LOXL2 expression in conditioned cell culture media and cell-derived ECM was analyzed by Western blotting.

For the role of LOXL2’s catalytic function on cell alignment in response to hypertensive mechanical loading, four conditions were examined: 1) HASMC controls, 2) T1 (LOXL2 depleted by CRISPR-Cas9 gene editing as previously described^38^; loss of function), 3) T1+ LOXL2 (recovery of function), in which LOXL2-enriched media, prepared by adenoviral overexpression of LOXL2 in T1 cells was used, and 4) HASMC + PAT-1251 (100 μM), a small molecule inhibitor of LOXL2 to pharmacologically inhibit LOXL2. Conditioned media was used to examine LOXL2 expression by Western blotting. Cell alignment analysis was performed as described next.

### Cell alignment analysis

PDMS membranes subjected to stretch (0, 10, 20%) were fixed in 4% formaldehyde and stained with Alexa Fluor 488 Phalloidin (ThermoFisher) to visualize actin. Images were captured with a confocal microscope (Leica SP8) at 5x magnification and imported into CurveAlign V4 software (https://github.com/uw-loci/curvelets/) run on MATLAB (2021 edition, The MathWorks, Natick, MA) to determine fiber orientation using a fast discrete curvelet transform-based algorithm. The curvelet orientations and the associated alignment coefficient, kurtosis, standard deviation, and variance parameters were obtained using a boundary-free CT fiber analysis method and a 0.06 fraction of kept coefficients. Intensity of fibers in the same orientation is characterized by the alignment coefficient which ranges from 0 to 1 with 1 signifying completely aligned fibers. Cumulative compass plots of curvelet orientation for each condition were generated using a custom MATLAB script. Plots were normalized to the number of replicates and plotted with a bin size of 5 degrees.

### Western blotting

Tissue specimens were homogenized, and cells were lysed in mammalian protein extraction reagent (M-PER; Thermo Fisher) containing protease inhibitors cocktail (Roche). Soluble proteins and insoluble matrix were separated by centrifugation at 10,000 x *g* for 15 minutes. A Bio-Rad protein assay was used to determine protein concentration in the soluble fraction. The insoluble matrix samples were resuspended in in 1.5x Laemmli buffer with DTT (New England Biolabs) in a normalized volume based on cytosolic protein concentration (50 μl per mg of soluble protein). Western blotting of tissue samples was performed using 25 µg of the soluble fraction, and 25 μl of the insoluble matrix. Secreted protein in cell culture media was enriched using StrataClean Resin (Agilent, 10 µl per 5 ml of media) as previously described^37, 42^.

Protein samples were resolved by SDS-PAGE and electro-transferred onto nitrocellulose membrane. Blots were stained with Ponceau S, imaged (BioRad ChemiDoc) and then blocked in 3% nonfat milk in TBST (Tris-buffered saline, 0.1% Tween 20). Blots were then incubated with primary antibody (1:1000, 2 h) washed and incubated with secondary antibody (1:10,000, 2 h). After 3 TBS-T washes, blots were developed with the Clarity Western ECL system (Bio-Rad). The following primary antibodies were used: LOX rabbit polyclonal (ThermoFisher PA1-46020), LOXL2 rabbit monoclonal (Abcam ab179810), collagen Type I Polyclonal Antibody (Thermo Scientific 14695-1-AP), COLIV rabbit polyclonal (Assay Biotech C0157), beta Actin Loading Control Monoclonal Antibody (Invitrogen, MA5-15739), GAPDH mouse monoclonal (Novus Bio NB300221). Primary antibodies were validated using knockdown/knockout (LOX, LOXL2), using recombinant protein (Collagens), or through prior reports in the literature (actin, GAPDH). Secondary antibodies used were goat anti-mouse IgG (H+L)-HRP conjugate (Biorad 1706516), AffiniPure goat anti-rabbit IgG (H+L)-HRP conjugate (Jackson ImmunoResearch 111035144).

### Histology

Fixed aortic samples were paraffin embedded and sectioned at 6 μm thickness with visible lumen and mounted onto charged slides (Reference Pathology Lab, JHU). Hematoxylin and eosin, Masson’s trichrome, and Movat pentachrome stains were performed using standard methods and imaged with a Laxco microscope at x4, x10, x20 and x40 magnification. Lumen diameter, aortic wall thickness, intralamellar distance, and VSMC nuclei in the medial layer were determined using ImageJ software.

### Statistical Analysis

Data are presented as mean ± standard error of the mean (SEM). Sample size (n) is indicated for each reported value. Persons performing statistical analyses were blinded to the groups, where appropriate. Shapiro-Wilk normality test and Kolmogorov-Smirnov normality test were used to test for and verify Gaussian distribution of data. Two means were compared using Student’s t-test and more than two means were compared by 1-way ANOVA with Tukey or Bonferroni *post hoc* analysis, as indicated in figure legends. For multiple (grouped) comparisons, 2-way ANOVA with Tukey or Bonferroni *post hoc* analysis was used, as indicated in figure legends. Means were considered to be statistically different at *P* < 0.05.

## Results

### LOXL2 depletion delays onset of Ang II induced hypertension in male mice

Systolic, diastolic, and mean blood pressure (BP) were measured in conscious mice immediately before (baseline) and during Ang II infusion weekly for a period of 3 weeks (**Figure 1A**). Baseline BP values were similar across all groups (**Figure 1B, C, D; Supplemental Figure S1**). Systolic BP increased rapidly in WT males with Ang II infusion, with a statistically significant increase within 1 week, and remained elevated for the duration of the study period. In the *Loxl2^+/-^* males with Ang II treatment, the onset of hypertension was delayed compared to WT, with a lower SBP at 1 week and reaching statistical significance at 2 weeks following Ang II infusion (**Figure 1B; Supplemental Figure S1**).

### Female mice exhibit delayed onset of hypertension when compared to males

BP elevation with Ang II infusion was slower in WT females than in WT male counterparts (**Figure 1B, C, D**), and similar to that noted in the Loxl2+/-males with Ang II (**Figure 1B, D**). A steady increase in BP was observed in both WT and *Loxl2^+/-^* females with Ang II infusion over the course of 3 weeks **(Figure 1C, D Supplemental Figure S1).** Magnitude of BP elevation in females was similar to WT males by 2 weeks of Ang II infusion.

### LOXL2 depletion prevents Ang II induced weight loss in both sexes and cardiac hypertrophy in males but not females

We evaluated the effect of Ang II induced hypertension on body weight **(Figure 1E),** heart weight **(Figure 1F)** and heart weight/body weight ratio **(Figure 1G)** as an index of cardiac hypertrophy. Consistent with prior studies^54–56^, Ang II infusion resulted in a significantly lower body weight in WT males (vs. control WT males) **(Figure 1E)**. *Loxl2^+/-^* control males had significantly lower body weight compared to WT controls and Ang II infusion did not result in lower body weight in *Loxl2^+/-^* males (vs. control *Loxl2^+/-^* males) (**Figure 1E**). Heart weight increased with Ang II infusion in both WT and *Loxl2^+/-^* males **(Figure 1F)**. Cardiac hypertrophy (increased heart weight to body weight ratio) was observed with Ang II infusion in both WT and *Loxl2^+/-^* males when compared to normotensive counterparts (**Figure 1E**).

Female WT and *Loxl2^+/-^* mice were lighter than their male counterparts (**Figure 1E)**; body weight was similar amongst female groups **(Figure 1E)**. Heart weight increased with Ang II infusion in both WT and *Loxl2^+/-^* females. Surprisingly, cardiac hypertrophy with Ang II induced hypertension was significantly higher in *Loxl2^+/-^* females but not in WT females (**Figure 1G**).

### LOXL2 depletion prevents elevation in PWV caused by Ang II-induced hypertension in males

PWV increased steadily in male WT mice over 3 weeks of Ang II infusion (**Figure 2A; Supplemental Figure S2).** Increase in BP **(Figure 1B)** preceded the onset of significant PWV increase **(Figure 2A)**. In *Loxl2^+/-^* males, PWV was not significantly elevated **(Figure 2A)** despite the eventual development of hypertension with Ang II infusion (**Figure 1B**).

**Figure 2.**
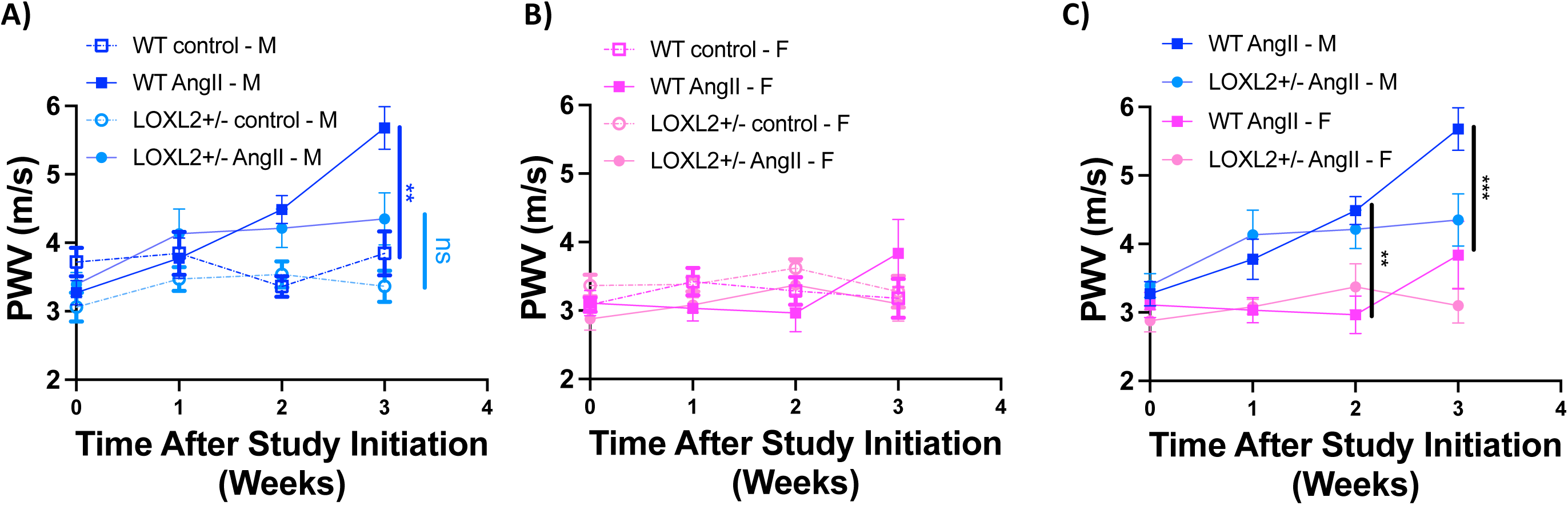
In vivo arterial stiffness assessment with Ang II induced hypertension in WT and *Loxl2^+/-^* male and female mice. **(A)** Pulse wave velocity (PWV) in male (M) WT and *Loxl2^+/-^* mice with and without Ang II infusion (n = 8 for WT control M, n = 8 for WT+Ang II M, n=8 for *Loxl2^+/-^* control M, and n = 7 for *Loxl2^+/-^* +Ang II M. At 3 wks: \*\**p* < 0.01 for WT Ang II M vs. WT control M, by repeated measures 2-way ANOVA with Tukey *post hoc* analysis; **(B)** PWV in female (F) WT and *Loxl2^+/-^* mice with and without Ang II infusion (n = 10 WT control F, n = 8 WT+Ang II F, n=8 *Loxl2^+/-^* control F, n = 8 and *Loxl2^+/-^* +Ang II F; no significance found by repeated measures 2-Way ANOVA with Tukey *post hoc* analysis) **(C)** PWV in male (M) and female (F) WT and Loxl2+/-mice with Ang II infusion (n = 8 for WT control M, n = 8 for WT+Ang II M, n=8 for *Loxl2^+/-^* control M, and n = 7 for *Loxl2^+/-^* +Ang II M, n = 10 WT control F, n = 8 WT+Ang II F, n=8 *Loxl2^+/-^* control F, n = 8 and *Loxl2^+/-^* +Ang II F; at 2 wks: **p<0.01 WT+Ang II M vs WT+Ang II F; at 3 wks: ***p<0.001 WT+Ang II M vs WT+Ang II F by 2-Way ANOVA with Sidak post hoc analysis).

### Female mice are protected from hypertension induced increase in PWV

In female mice, Ang II infusion did not lead to elevated PWV in either WT or *Loxl2^+/-^* groups (**Figure 2B, C**) despite the increase in systolic BP (**Figure 1C**). Heart rate was similar in all groups during PWV measurements (**Supplemental Figure S2**). WT females with Ang II infusion had markedly lower PWV than male counterparts by 2 weeks of Ang II infusion (**Figure 2C**).

### LOXL2 depletion protects males from Ang II mediated potentiation of vasoconstriction

We next examined the changes in vascular reactivity responses arising from Ang II-induced hypertension by wire myography. A marked increase in maximal vasoconstriction response to phenylephrine (PE) was observed in Ang II-treated WT males, but not in *Loxl2^+/-^* males (**Figure 3Ai**). EC50s for PE-induced constriction were similar in all groups of male mice **(Figure 3Bi)**. Acetylcholine-induced endothelial-dependent vasorelaxation was significantly impaired with Ang II-induced hypertension in WT males with attenuated relaxation and higher EC50. LOXL2 depletion was partially protective with improvements in maximal relaxation and EC50 in *Loxl2^+/-^* hypertensive males (**Figure 3Aii, Bii**). Endothelial independent responses to sodium nitroprusside were similar in males of all groups (**Figure 3Aiii, Biii**).

**Figure 3.**
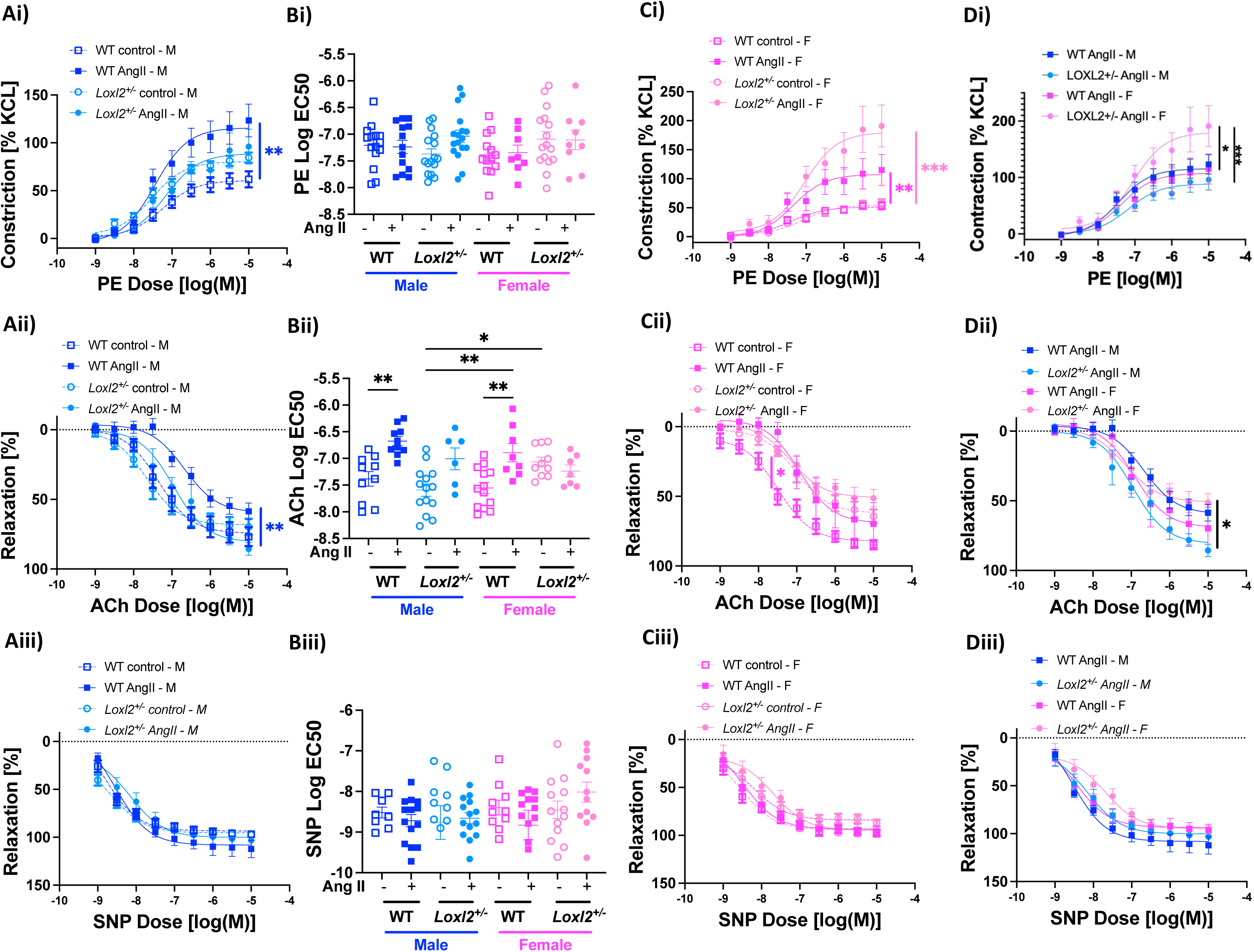
Effect of sex and LOXL2 depletion on hypertension induced vascular dysfunction. Analysis of aortic rings from male and female mice to increasing concentrations of phenylephrine (PE; **Ai:** male, **Bi:** EC50, **Ci:** female, **Di:** Male vs. female hypertensives), calculated as percentage of max KCl-induced vasoconstriction, Acetylcholine (Ach) - induced endothelium-dependent vasorelaxation of vessels pre-constricted by PE (**Aii:** male, **Bii:** EC50, **Cii:** female, **Dii:** Male vs. female hypertensives), and Endothelium-independent vasorelaxation in response of PE pre-constricted rings to increasing concentrations of sodium nitroprusside (SNP) (**Aiii:** male, **Biii:** EC50, **Ciii:** female, **Diii:** Male vs. female hypertensives) (*n* = 7 – 10 mice per cohort; two rings were assessed from each animal. \**p* < 0.05, \*\**p* < 0.01, \*\*\**p* < 0.001; ### p<0.01 Female Loxl2+/- hypertensive vs. all other groups in in panel **Di**, by repeated measures 2-Way ANOVA with Bonferroni post hoc analysis).

### LOXL2 depletion worsens vascular contractility in female hypertensives

In females, maximal contractility to phenylephrine was elevated with Ang II infusion in WT mice and to a larger magnitude in *Loxl2^+/-^* mice compared to corresponding controls (**Figure 3Ci**) and when compared with hypertensive males (**Figure 3Di**). EC50s for PE-induced constriction were similar amongst all groups of female mice (**Figure 3Bi**) and were similar to males. Endothelial dependent relaxation in response to increasing concentrations of acetylcholine was impaired with Ang II in WT female mice when compared with normotensive WT females, with an increase in EC50 (**Figure 3Bii);** Loxl2+/- females did not exhibit worsening endothelial dependent relaxation with Ang II induced hypertension. Surprisingly however, normotensive *Loxl2^+/-^* females exhibited impaired relaxation response to acetylcholine when compared with *Loxl2^+/-^* males, with higher EC50 (**Figure 3Bii, Dii**). Endothelial independent vasorelaxation elicited by sodium nitroprusside was similar in all groups **(Figure 3Biii, Ciii, Diii)**.

### LOXL2 depletion attenuates passive arterial stiffening in males

Hypertension resulted in a marked increase in passive stiffness of intact aortic rings from WT males (**Figure 4Ai)** that was attenuated in *Loxl2^+/-^* males **(Figure 4Bi)**. Incremental moduli at strains of 0.5 (elastin deformation) and 2.5 (collagen deformation) were also markedly elevated in intact rings from WT males with Ang II infusion **(Figure 4Ci, Cii).** Decellularized aortic rings from male WT mice that received Ang II were also stiffer than untreated counterparts (**Figure 4Ei**) and *Loxl2^+/-^* males had an attenuated response (**Figure 4Fi**). The incremental moduli of decellularized aortae from hypertensive WT males were similar at low (0.5) strain and significantly higher at strain of 2.5 (**Figure 4Gi, ii**), suggesting that collagen accumulation is a key component of aortic ECM stiffening. Interestingly, Ang II induced hypertension resulted in elevated ultimate tensile strength (stress at failure) in the *Loxl2^+/-^* males in both the intact and decellularized aorta, but not in WT mice (**Figure 4Di, Hi**). Strain at failure was similar in the intact rings from males but was increased with Ang II infusion in decellularized rings from both WT and *Loxl2^+/-^* males **(Figure 4Dii, Hii).**

**Figure 4.**
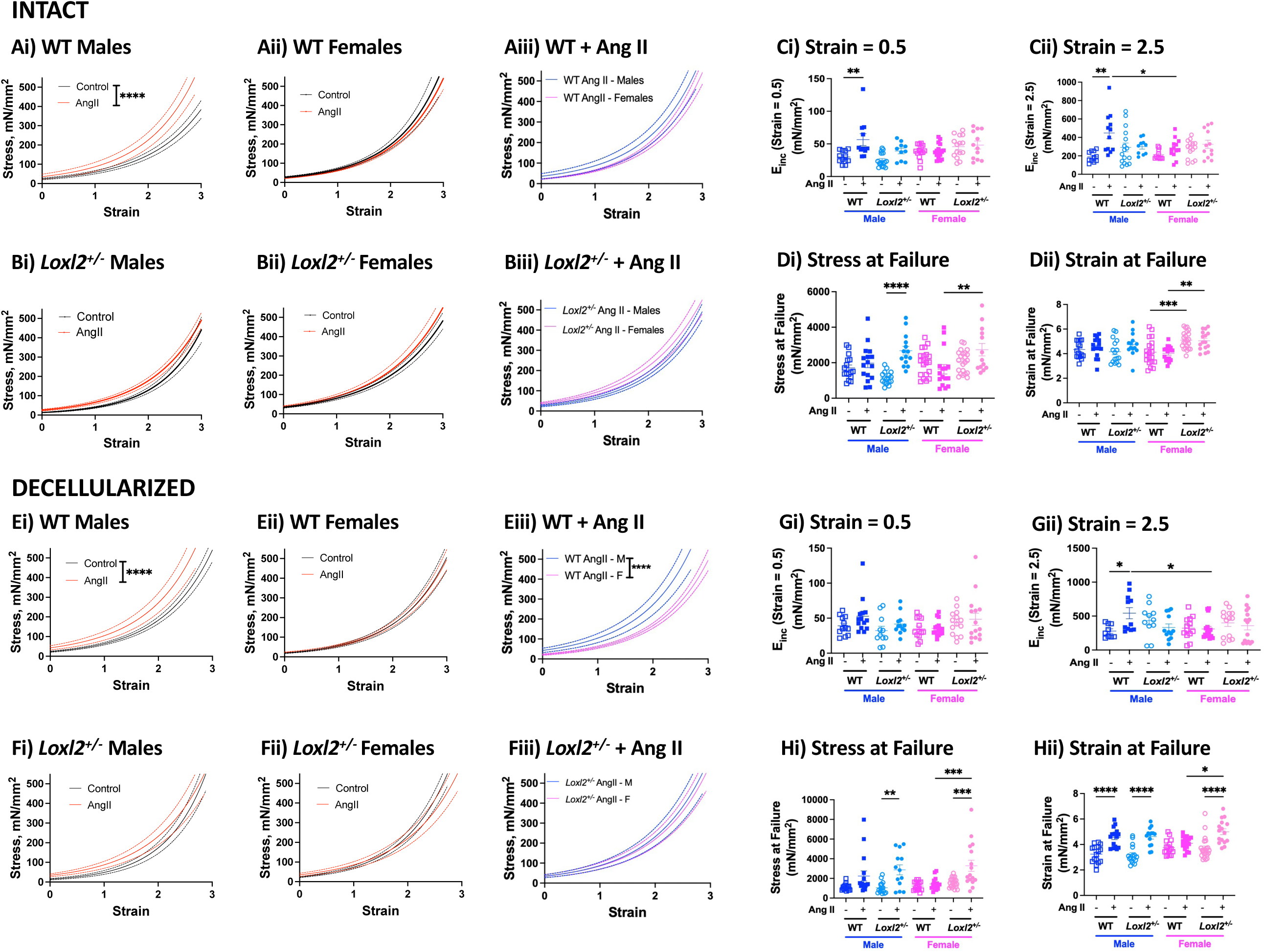
Effects of Ang II infusion on passive mechanical properties of the aorta in male and female WT and *Loxl2^+/-^* mice. Uniaxial tensile testing of **(A-D)** intact and **(E-H)** decellularized aortic segments from male and female WT and *Loxl2^+/-^* mice with and without Ang II infusion. **(A, B)** Stress-strain curves of intact aortic rings from WT (**Ai:** male hypertensive vs. normotensive, **Aii:** female hypertensive vs normotensive, **Aiii:** hypertensive male vs. female) and *Loxl2^+/-^* mice (**Bi:** male hypertensive vs. normotensive, **Bii:** female hypertensive vs normotensive, **Biii:** hypertensive male vs. female). Data are shown as mean (solid line)L±Lstandard error (dotted lines). **(C)** Incremental elastic modulus of intact aortic segments at **(Ci)** low strain (0.5) in and **(Cii)** high strain (2.5). **(D)** Ultimate stress **(Di)** and strain **(Dii)** at failure of intact aortic rings. **(E)** Stress-strain curves of decellularized aortic rings from WT (**Ei:** male hypertensive vs. normotensive, **Eii:** female hypertensive vs normotensive, **Eiii:** hypertensive male vs. female) and *Loxl2^+/-^* mice (**Fi:** male hypertensive vs. normotensive, **Fii:** female hypertensive vs normotensive, **Fiii:** hypertensive male vs. female). Data are shown as mean (solid line)L±Lstandard error (dotted lines). **(F)** Incremental elastic modulus of decellularized aortic segments at **(Fi)** low strain (0.5) in and **(Fii)** high strain (2.5). **(G)** Ultimate stress **(Gi)** and strain **(Gii)** at failure of decellularized aortic rings. (n = 14-22 aortic rings; 2 rings were tested from n = 7-11 mice in each group; ****p<0.0001 by repeated measures 2-way ANOVA with Bonferroni post-hoc analysis for stress vs strain curves; *p<0.05, **p<0.01, ***p<0.001, ****p<0.0001 by 2- way ANOVA with Bonferroni post-hoc analysis for scatter plots).

### WT females are protected from hypertension induced aortic remodeling and LOXL2 depletion improves aortic mechanics in hypertensive females

Intact aortae of both WT and *Loxl2^+/-^* females were protected from stiffening in response to Ang II-induced hypertension and exhibited no change in incremental elastic modulus at low (0.5) or high (2.5) strain **(Figure 4Aii, Bii, Ci, Cii)**. Similarly, decellularized aortae of females did not show any change in material properties with Ang II infusion **(Figure 4Eii, Fii, Gi, Gii)**. Intact aortae of hypertensive WT females were modestly more compliant than WT hypertensive males and exhibited lower elastic modulus at high (2.5) strain (**Figure 4Aiii, Cii**). On the other hand, decellularized aortae from hypertensive WT females were markedly more compliant than those from hypertensive WT males **(Figure 4Eiii, Gii)**. This difference between the sexes was abolished in the *Loxl2^+/-^* female mice **(Figure 4Biii, Cii, Fiii, Gii)**.

In females, yield stress of intact rings remained unchanged in both genotypes with Ang II infusion when compared with their normotensive counterparts (**Figure 4Di**). Interestingly, however, hypertensive *Loxl2^+/-^* females exhibited higher yield stress than hypertensive WT females **(Figure 4Di)**. Normotensive and hypertensive *Loxl2^+/-^* females exhibited higher strain at failure when compared with WT counterparts **(Figure 4Dii**). Decellularized rings of *Loxl2^+/-^* females with Ang II had higher yield stress when compared to both normotensive *Loxl2^+/-^* females and hypertensive WT females (**Figure 4Hii**). Strain at failure was higher in the *Loxl2^+/-^* females vs. WT littermate females and was unaffected by Ang II (**Figure 4Hi**). Decellularized segments from *Loxl2^+/-^* females exhibited higher strain at failure than both the WT females with Ang II as well as *Loxl2^+/-^* female controls (**Figure 4Hii**).

### LOXL2 depletion attenuates aortic wall remodeling caused by Ang II-induced hypertension in males

We next examined whether aortic remodeling was LOXL2 dependent using Hematoxylin Eosin (H&E), Masson’s trichrome, and Movat’s pentachrome staining **(Figure 5Ai)**. Ang II infusion resulted in VSMC hyperplasia in WT males, reflected in the increased VSMC nuclei density in the vascular wall, but not in the *Loxl2^+/-^* males (**Figure 5B).** Aortic lumen diameter was similar amongst all groups in males (**Figure 5C**), but wall thickness increased with Ang II infusion in both WT and *Loxl2^+/-^* males **(Figure 5D).** Intralamellar distance was higher in both WT and *Loxl2^+/-^* males with Ang II infusion when compared to normotensive counterparts (**Figure 5E**).

**Figure 5.**
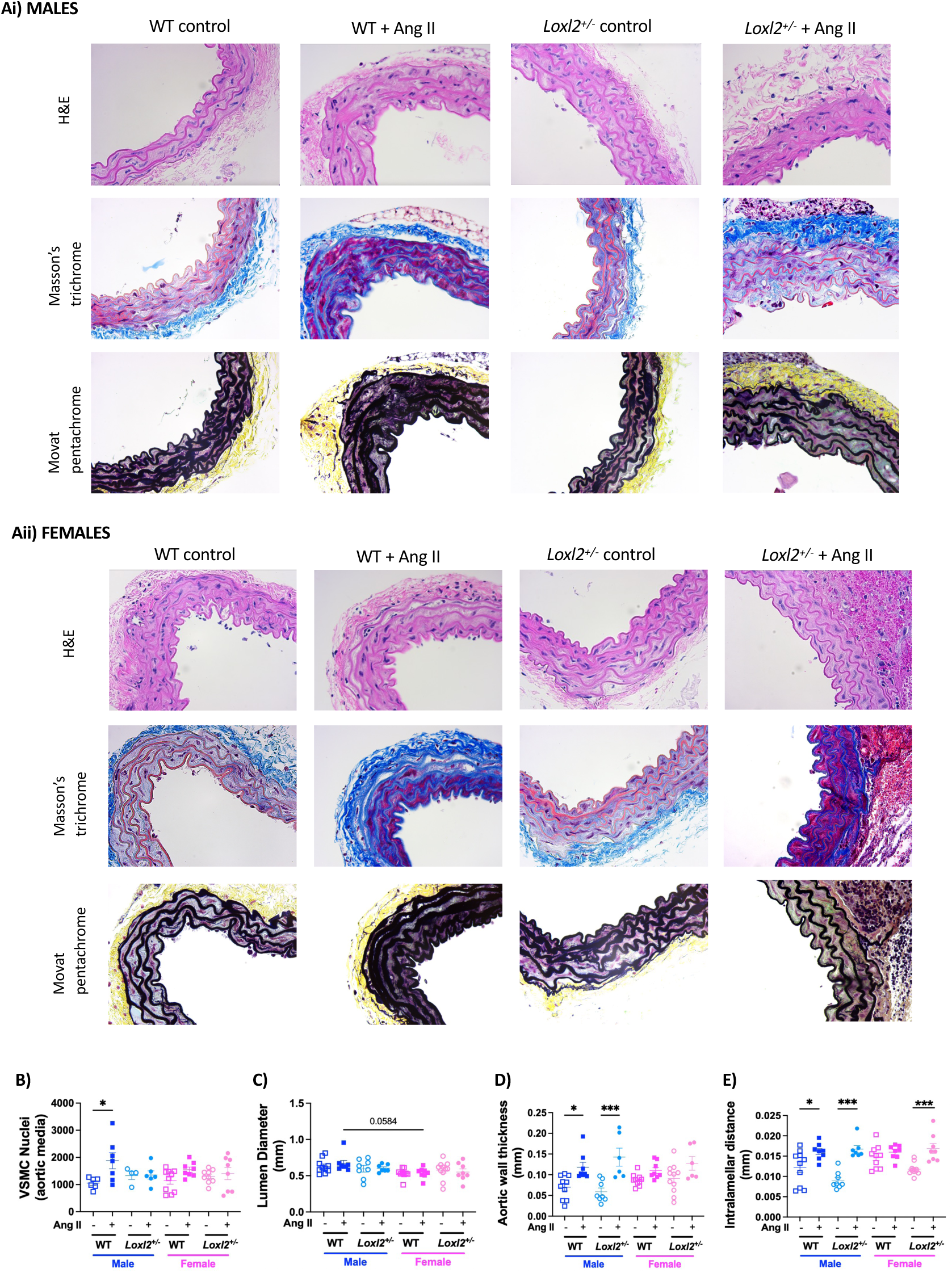
Histochemical analysis of aorta from WT and *Loxl2^+/-^* mice. **(A)** Representative Hematoxylin and Eosin (H&E), Masson’s trichrome, and Movat pentachrome stains in **(Ai)** male and **(Aii)** female mice. (**B)** VSMC nuclei in vascular wall **(C)** Lumen diameter, **(D)** wall thickness and **(E)** intralamellar distance calculated from H&E stained images. (*n* = 5-7 for each group, \*\**p* < 0.01, and \*\*\**p* < 0.001 by 2-way ANOVA with Bonferroni post hoc analysis).

### Females are protected from Ang II induced wall remodeling

Hypertensive WT female mice showed minimal VSMC hyperplasia when compared with normotensive WT females and hypertensive WT males (**Figure 5 Aii, B**). Aortic lumen diameter was similar amongst all groups in females, and hypertensive WT females exhibited a strong trend towards lower lumen diameter than hypertensive WT males (**Figure 5C**). Wall thickness was similar in all groups in the female mice and similar to males **(Figure 5D).** Normotensive WT females exhibited a strong trend towards higher intralamellar distance than the normotensive *Loxl2^+/-^* female littermates. Intralamellar distance increased in the female hypertensive *Loxl2^+/-^* group but not in female WTs when compared with their normotensive counterparts (**Figure 5Eii**).

### LOXL2 accumulation and processing are increased with an increase in collagen I and decrease in collagen IV in the aortic ECM of hypertensives of both sexes

Male and female aorta were examined by Western blotting on separate gels (**Figure 6Ai, Aii**). To obtain insights on LOX/LOXL2 function in the vessel wall, we evaluated full length and processed LOX/LOXL2, collagen I, and collagen IV protein abundance in the decellularized aortic ECM by Western blotting. Both full length and processed LOXL2 were elevated in WT males with Ang II infusion (**Figure 6Ai, Bi, Bii**). In the *Loxl2^+/-^* mice, although a strong trend towards increased levels of both full length and processed LOXL2 were noted with Ang II infusion, the data did not reach statistical significance (**Figure 6Ai, Bi, Bii**). In males, pro-LOX levels were similar in all groups and a trend towards increased active LOX was noted in the Ang II treated WT and *Loxl2^+/-^* groups (**Figure 6Ai, Biii, Biv**). In males, collagen I increased, and collagen IV decreased in the aortic ECM with Ang II treatment (**Figure 6Ai, Bv, Bvi**).

**Figure 6.**
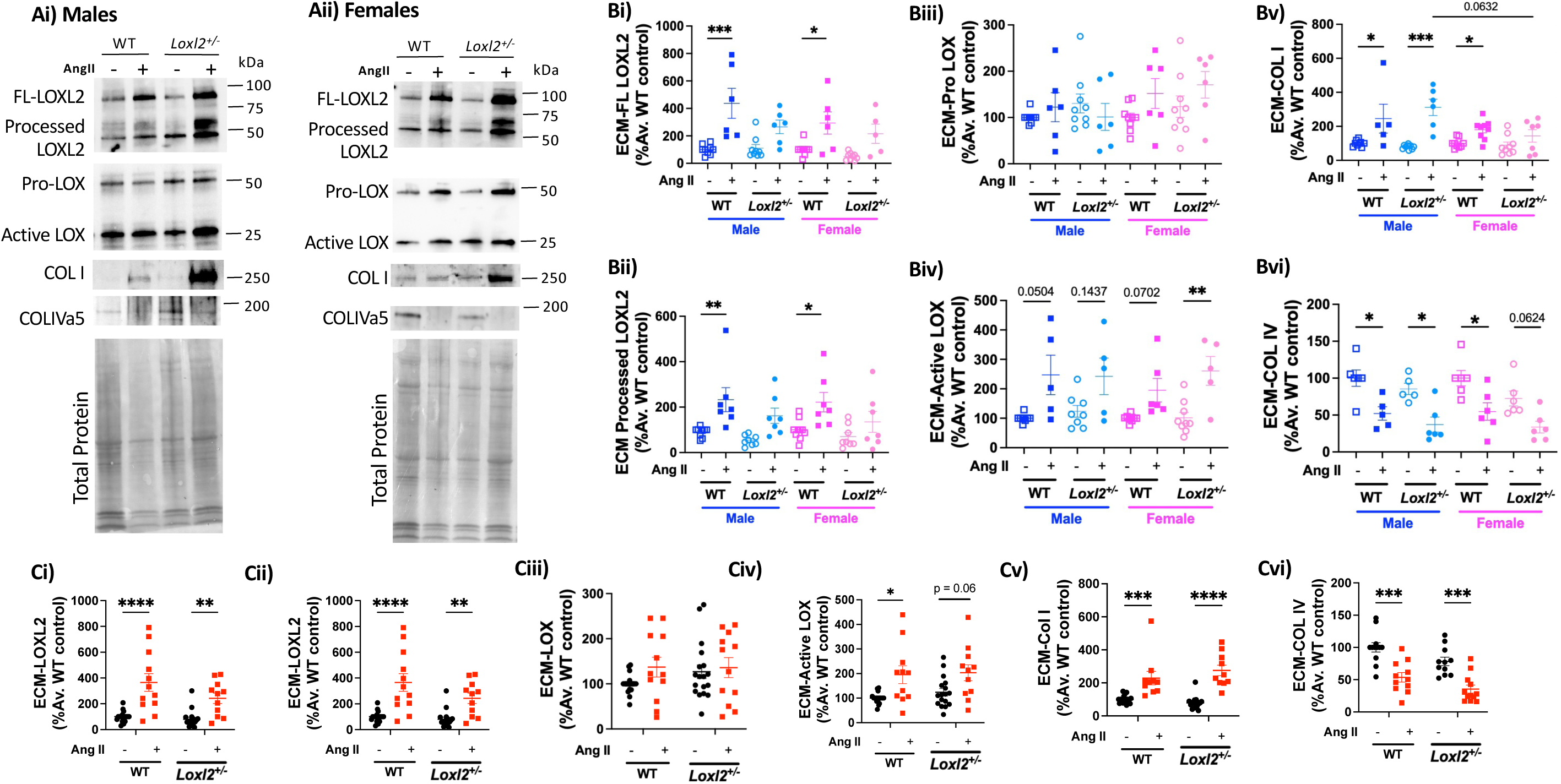
Assessment of LOX, LOXL2, collagen I, and collagen IV protein expression in aortic ECM by Western blotting. **(A)** Representative Western blots showing expression of LOXL2, LOX and COL I in the extracellular matrix (ECM), fraction of aorta extracted 3 wks after initiation of experiment (**Ai:** males, **Aii:** females). **(B)** Sex-disaggregated densitometry analyses of **(Bi)** full length LOXL2, **(Bii)** processed C-terminus fragment of LOXL2, **(Biii)** pro-LOX, **(Biv)** processed, active LOX, **(Bv)** collagen I, and **(Bvi)** collagen IV. (n = 5-9 mice per group; \**p* < 0.05, \*\**p* < 0.01, and \*\*\**p* < 0.001 by 2-way ANOVA with Tukey *post-hoc* analysis). **(C)** Combined densitometry analyses of **(Ci)** full length LOXL2, **(Cii)** processed C-terminus fragment of LOXL2, **(Ciii)** pro-LOX, **(Civ)** processed, active LOX, **(Cv)** collagen I, and **(Cvi)** collagen IV. (n = 10-18 mice per group; \**p* < 0.05, \*\**p* < 0.01, and \*\*\**p* < 0.001, ****p<0.0001 by 2-way ANOVA with Tukey *post hoc* analysis).

In females, full length and processed LOXL2 significantly increased with Ang II treatment in WT females but not in *Loxl2^+/-^* females although a strong trend was noted (**Figure 6Aii, Bi, Bii).** Pro-LOX was similar amongst all female groups, and active LOX was elevated in *Loxl2^+/-^* females with Ang II infusion (**Figure 6Aii, Biii, Biv).** LOX and LOXL2 levels were similar to male counterparts (genotype and Ang II matched; **Figure 6Ai, ii; Bi-iv**). Collagen I was markedly higher, and collagen IV was lower with Ang II infusion in hypertensive WT females when compared with normotensive WT females (**Figure 6Aii, Bv, Bvi**). Hypertensive *Loxl2^+/-^* females exhibited similar collagen I levels but reduced collagen IV abundance when compared with normotensive *Loxl2^+/-^* females (**Figure 6Aii, Bv, Bvi)**. Hypertensive WT and *Loxl2^+/-^* mice exhibited a strong trend towards lower collagen I than hypertensive male counterparts (**Figure 6Bv, vi**).

When the sexes were grouped together to increase statistical power, we noted increased protein levels of full length and processed LOXL2 in both WT and *Loxl2^+/-^* mice with Ang II infusion versus untreated counterparts (**Figure 6Ci, ii**). While full length LOX was similar in all groups, active LOX was elevated in the hypertensive WTs when compared with normotensive WTs, and a strong trend was noted in the *Loxl2^+/-^* hypertensives when compared with *Loxl2^+/-^* normotensive group **(Figure 6Ciii, iv)**. Hypertension led to elevated ECM collagen I and reduced collagen IV in both WT and *Loxl2^+/-^* mice **(Figure 6Cv, vi)**.

### Cyclic strain upregulates LOXL2 expression in VSMCs but not ECs

We next examined the regulation of LOXL2 protein expression by normotensive (10%) versus hypertensive (20%) cyclic biomechanical strain in HAECs and HASMCs. Both HASMCs and HAECs aligned perpendicularly to the direction of stretch (**Figure 7Ai, Aii**). In HASMCs, cyclic strain had no impact on LOXL2 abundance in the conditioned cell culture media (**Figure 7Bi, Bii**). In the cell-derived ECM, while full length LOXL2 was similar in all groups, abundance of processed LOXL2 was markedly elevated with increasing strain (**Figure 7Bi, iii, iv**). In HAECs, LOXL2 level was highest at static state, and decreased significantly at both 10% and 20% stretch (**Figure 7Ci-iii)**. Thus, these results point to the VSMCs as the primary source of LOXL2 in the vascular media.

**Figure 7.**
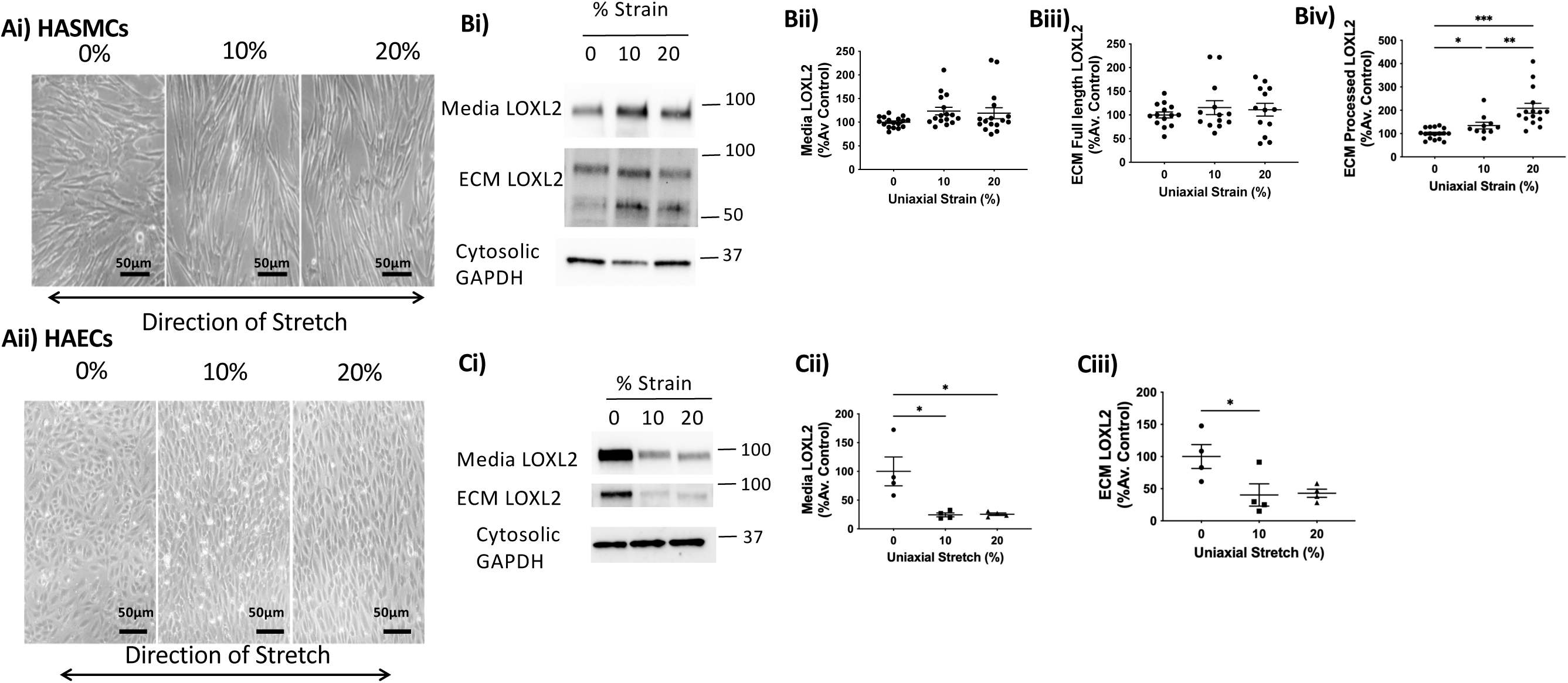
Processed LOXL2 abundance increased with cyclic strain in VSMCs but not ECs. **(A)** representative phase contrast images of **(Ai)** human aortic smooth muscle cell (HASMCs) and **(Aii)** human aortic endothelial cells (HAECs) under 0% (static condition), 10%, and 20% cyclic strain. Scale bar = 50 μm. Double-headed arrow indicates the strain vector. **(B, C)** Representative Western blots and densitometry analysis of LOXL2 expression in **(B)** HASMC (**Bi:** representative blots, **Bii:** LOXL2 in conditioned cell culture media, **Biii:** Full length LOXL2 in cell-derived ECM, **Biv:** Processed LOXL2 in cell derived ECM; n = 13 per group) and **(C)** HAEC (**Ci:** Representative blots **Cii:** LOXL2 in conditioned cell culture media, **Biii:** Full length LOXL2 in cell-derived ECM; n = 4 per group). (\**p* < 0.05, \*\**p* < 0.01, ***p<0.001 by ordinary 1-way ANOVA with Bonferroni post hoc analysis.).

### LOXL2 plays an important role in VSMC alignment

We next determined if LOXL2 facilitates VSMC alignment when exposed to stretch through catalytically dependent or catalytically independent mechanisms. To this end, we compared the realignment of HASMCs in response to a hypertensive strain of 20% for 24 hours, with and without LOXL2’s crosslinking function. Four groups were compared: 1) WT HASMC, 2) WT HASMC + PAT-1251, a specific inhibitor of LOXL2’s catalytic function, 3) T1 HASMCs in which *Loxl2* gene was knocked-out by CRISPR-Cas9 gene editing (loss of function), and 4) T1 HASMCs + LOXL2 (recovery of function) to reintroduce LOXL2. At the end of the uniaxial stretching period, LOXL2 abundance in the culture media determined by Western blotting confirmed loss of LOXL2 expression in the T1-HASMCs and recovery of LOXL2 expression in the T1+LOXL2 condition (**Figure 8Ai, Aii**). Confocal microscopy of FITC-Phalloidin labeled cells revealed that WT HASMCs oriented orthogonal to the stretch vector, as evidenced by the increase in alignment coefficient in 20% strain versus static (0%) condition **(Figure 8Bi, Bii, Biii)**. Both LOXL2 inhibition (PAT-1251) and knockout (T1 HASMC *LOXL2^-/-^* cells) resulted in loss of cyclic strain induced alignment of HASMCs, evidenced by the loss/diminished change in alignment coefficient between the static and uniaxial strain conditions (**Figure 8Bi, Bii, Biii**). Importantly, provision of LOXL2 restored the orientation response (i.e, increase in alignment coefficient upon exposure to uniaxial strain) in T1 cells. Together these findings suggest an important role for LOXL2 in mechanosensing and the involvement of its catalytic activity.

**Figure 8.**
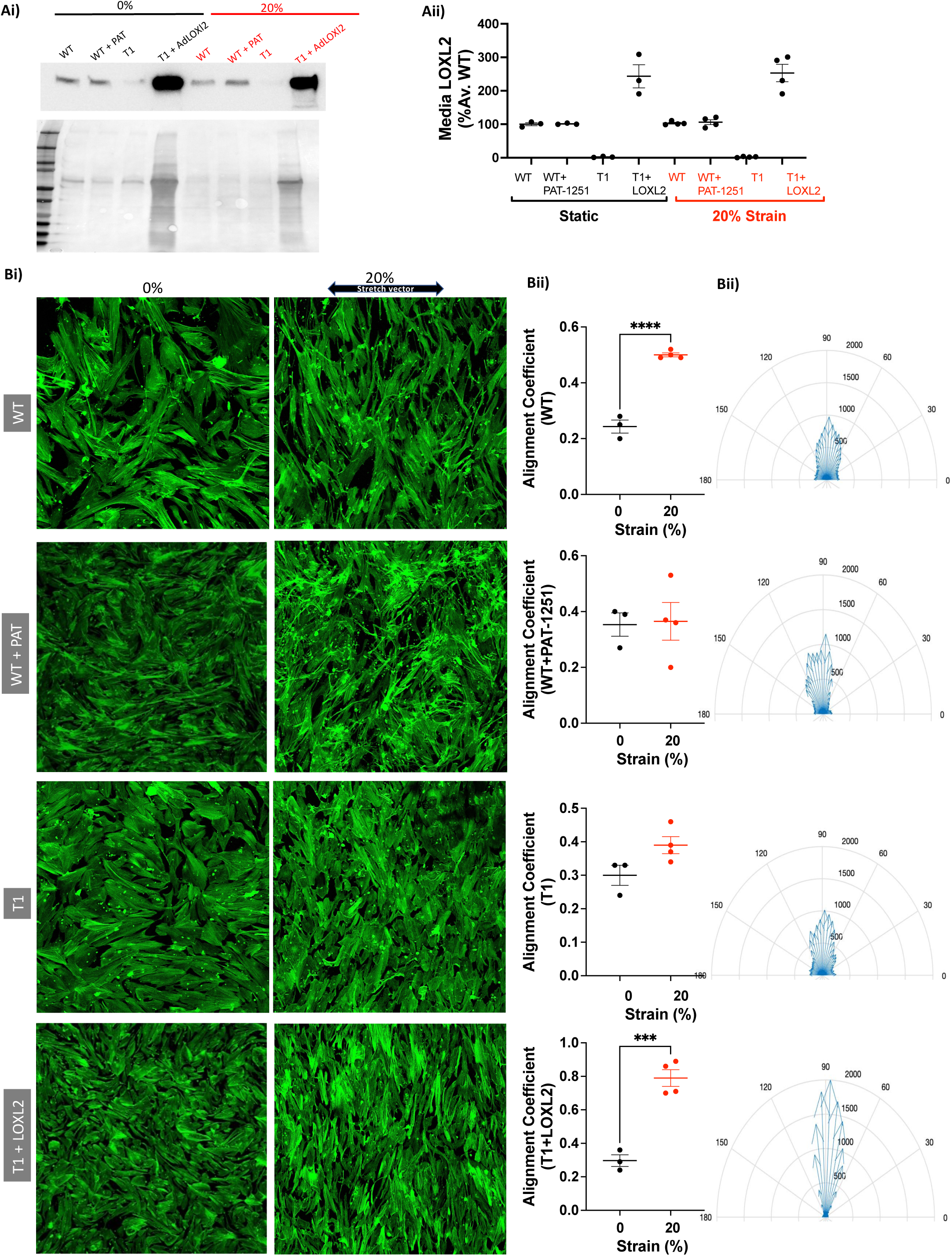
LOXL2 plays an important role in VSMC alignment. Wild-type (WT) HASMCs ± PAT-1251 (10μM) and LOXL2 knocked-out (T1) HASMCs ± Ad-LOXL2-enriched media were static or stretched at 20% on FN-coated PDMS chambers. **(Ai)** Representative Western blotting and **(Aii)** densitometry analysis of LOXL2 in conditioned cell culture media post 24-hr stretch. **(Bi)** Representative confocal microscopy images of formalin-fixed cells stained with FITC-phalloidin (green). Scale bar = 400 μm. **(Bii)** Alignment coefficient of cells assessed by orientation actin fibers by CurveAlign; **(Biii)** Representative cumulative compass plots showing the proportions of actin fibers at each angle. (Number of x5 magnification confocal images n = 3-4 per group, with 500+ cells per image, \*\*\**p* < 0.01, \*\*\*\**p* < 0.0001 by 2-way ANOVA).

## Discussion

In this study, we evaluated sex disparities in Ang II-induced hypertension in young adulthood, specifically, if differences in ECM remodeling are a major contributing factor in hypertension induced arterial stiffening in male and female mice. As LOXL2 is shown to mediate aortic stiffening in aging, we investigated whether central vascular stiffening in hypertension can be prevented by targeting LOXL2. The central findings were that development of hypertension, and the associated stiffening of conduit vessels occurs through distinct mechanisms and timelines in males and females during reproductive years when challenged with the same dose of Ang II. Effects of sex were prominent in WT mice including earlier onset of hypertension in males than in females and body weight loss in hypertensive males, but not in females. While males developed central arterial stiffening in vivo (indicated by higher PWV) following BP increase, female mice did not, despite onset of hypertension. When considered together with the literature showing that females are relatively protected from aging related arterial stiffening until menopause is reached^44^, these findings suggest a similar protection against stiffening in hypertension in young females. In this context, the effect of menopause and aging on the hypertension-induced aortic stiffening response in females remains to be studied. However, with the slower but steady increase in BP, it is also possible that later time points (>3 weeks) could lead to elevation in PWV, and this remains to be tested.

The viscoelastic nature of the compliance vessels means that hypertensive BP itself can lead to a higher PWV and ex vivo studies are needed to establish changes to inherent aortic mechanics. Our choice of studies were guided by the fact that PWV is the manifestation of the combined effects of active (VSMC tone) and passive (ECM stiffness) properties of the vessel We therefore examined if the elevation of PWV arises from vascular dysfunction and/or ECM remodeling by measuring vascular reactivity and passive aortic stiffness at the tissue level. Wire myography revealed sex-independent impairments in smooth muscle contractility with Ang II induced hypertension in WT mice, demonstrating that female sex is not protected from hypertension induced and VSMC dysfunction. Thus, improved vascular tone/function cannot fully account for the protection from PWV elevation in hypertensive females.

Endothelial function was impaired in WT male hypertensive aorta (versus normotensive WT male), as evidenced by impaired relaxation to acetylcholine without any change in sodium nitroprusside induced endothelial-independent relaxation. These shifts can be ascribed to impaired nitric oxide (NO) bioavailability, as the effects of the COX pathway were excluded using the COX inhibitor indomethacin. In WT females, while a shift in EC50 was noted, maximal relaxation was similar between hypertensive and normotensive females. This suggests that female sex is partially protected from endothelial dysfunction in hypertension. This could be due to estrogen, which is shown to play a role in protecting the vascular endothelium by increasing nitric oxide (NO) bioavailability through both the upregulation of endothelial NO synthase^13, 14^ and increase in intracellular free Ca^2+^ concentration in endothelial cells^15^.

Passive stiffness is defined by the combined effects of the ECM (composition and stiffness) and the VSMC (inherent stiffness) and the interactions between the two (e.g., strength and number of focal adhesions). The contributions of cells and ECM to overall stiffness were evaluated by comparing the mechanics of the intact aorta (=cells+ECM) and the decellularized aorta (=ECM alone), determined by tensile testing. In WT males, aortic stiffening in hypertension arose from both the cellular and ECM components. Specifically, in the hypertensive male aorta, VSMCs are important determinants of aortic modulus at low strain (0.5; elastin mediated deformation) and ECM remodeling governs the stiffness at high strains (2.5; collagen mediated deformation). This was further supported by overall changes in wall structure and composition including VSMC hyperplasia, increased wall thickness, increased intra-lamellar distance, and collagen accumulation in the male hypertensive aorta when compared with normotensive counterparts. The impairments in endothelial function identified by wire myography could, in part, underlie these changes, as loss of NO bioavailability is shown to induce aortic stiffening in aging.^57^

Female WTs were refractory to hypertension induced ECM remodeling, and thus preserved a more compliant ECM, stable wall structure, and stable PWV despite the onset of endothelial and smooth muscle dysfunction with hypertension. While the specific mechanisms that guide sex differences in ECM remodeling remains to be studied, the putative role of estrogen in mediating ECM remodeling emerges as an important consideration that is supported by the literature showing sexual dimorphisms in cardiovascular remodeling during aging.^58^ This is in part shown to be mediated by sex hormone receptors through regulation of fibrotic-specific ECM pathways^58^; for example, androgen signaling is associated with increased fibrotic processes. Collagen content and subtype are also shown to be modulated in a sex-specific manner. Female hearts were shown to contain more collagen I and III than men in youth, but higher collagen I and III with aging^59^. This is substantiated in our study as hypertensive young females had lower collagen I than hypertensive males. The specific effects of sex-hormone signaling and the differences between androgen and estrogen effects remains to be further studied and represents an important element of female health.

Given the substantive role of ECM remodeling in male and female aortic stiffening in hypertension, we focused on the putative role of LOXL2, an enzyme previously shown to contribute to age-associated aortic stiffening via regulation of VSMC tone/reactivity and ECM remodeling^37^. In vivo, in males, LOXL2 depletion delayed the onset of hypertension and prevented PWV elevation with hypertension whereas female sex was overall protective against PWV elevation independent of LOXL2. At the tissue level, in males LOXL2 depletion was clearly protective against hypertension induced aortic derangements, conferring improved contractile (VSMC) function and endothelial function. In addition, LOXL2 depletion improved the maximal load bearing capacity (higher yield stress and strain without any change in incremental elastic modulus (E_inc_)) of the aortic ECM in hypertensive males, indicating improved material properties. In females, LOXL2 depletion yielded benefits including prevention of hypertension induced endothelial dysfunction and a surprising improvement in load-bearing capacity even in normotensive mice (illustrated by a higher yield stress and yield strain without any change in E_inc_). However, in females, LOXL2 depletion also aggravated hypertension induced hypercontractility to phenylephrine. Thus, LOXL2 depletion can be considered as being detrimental for vascular reactivity but protective with regard to aortic ECM remodeling and mechanics in young female hypertensives. Taken together, these findings show that LOXL2 depletion ameliorates hypertension induced vascular matrix remodeling and improves load bearing in young males and females.

We next examined the vascular wall structure and composition to gain insights into the mechanisms by which LOXL2 confers benefits to mechanics of the hypertensive aorta and whether sex differences exist in this regard. Targeting LOXL2 prevented VSMC hyperplasia, indicative of de-differentiation and proliferation, in hypertensive males. Female sex was overall protective, with no change in wall thickness or VSMC content with hypertension, independent of LOXL2 depletion.

However, both male and female *Loxl2^+/-^* hypertensive mice still exhibited some evidence of structural alterations including an increase in intralamellar distance and a qualitative increase in collagen content identified by Masson’s trichrome and MOVAT pentachrome staining, suggesting the presence/activity of remodeling pathways. Western blotting revealed that both LOXL2 and LOX are involved in hypertensive aortic ECM remodeling in both sexes, as evidenced by the marked increase in full length and processed LOXL2 as well as active LOX in the aortic ECM of WT mice. Furthermore, consistent with prior studies that showed that LOXL2 processing shifts its substrate preference to collagen I^42^, and LOX exhibits preference for fibrillar collagens^60, 61^, collagen I content increased and collagen IV content decreased in the hypertensive aorta. While we only tested collagen I by Western blotting, the elevated levels of processed LOXL2 and active LOX suggest that other fibrillar collagens would accumulate in the hypertensive aorta, and this remains to be tested. The only sex difference noted in this regard was that female *Loxl2^+/-^* hypertensives induced a compensatory increase in prototypic LOX expression and the specific pathways and mechanisms underlying this remain to be elucidated. As both sexes exhibited similar shifts in LOX/LOXL2 and collagen I/IV abundance in the ECM, we combined the protein expression data from the two sexes to enhance statistical power to evaluate the effect of hypertension. This analysis shows that hypertension elicits LOXL2 and LOX activation combined with higher collagen I and lower collagen IV the aortic ECM, which explains the shifts in mechanical properties with hypertension. Moreover, the data show that a single allele of LOXL2 is sufficient to activate LOXL2 protein in the hypertensive aorta, although to a lower magnitude than in WT. This attenuated level of LOXL2 can still guide the shifts in collagen subtype content in the aortic ECM and is sufficient to preserve aortic mechanical stability in hypertension. Interestingly, while females exhibit this shift in collagen composition at the molecular level, they do not exhibit aortic stiffening as measured by tensile testing or PWV. Our data suggest that it is the attenuation of VSMC hyperplasia along with in vivo tone, defined by the net balance of vasoconstrictive and vasorelaxant milieu, that guide this benefit to overall tissue stiffness ex vivo and in vivo in females.

While in vivo experiments show the increased LOXL2 in the aortic ECM, they do not define the cellular source of LOXL2 or whether biomechanical strain regulates LOXL2. *Ex vivo* experiments with HASMCs and HAECs revealed that VSMCs are a more significant source of LOXL2 under normotensive and hypertensive cyclic strain imposed by the cardiac cycle, but ECs are not. Furthermore, the increase in processed LOXL2 in the HASMC-derived ECM with hypertensive loading as well as in the aortic ECM of hypertensive mice suggests that LOXL2 processing in the hypertensive aorta can arise due to mechanical loading/ mechanosensing. On the other hand, full length LOXL2 is unchanged in cells exposed to hypertensive strain but is higher in the hypertensive aorta. This suggests that elevated full length LOXL2 in the hypertensive aorta cannot be explained by mechanical loading and involves other pathways/ mechanisms. Prior studies show that loss of NO bioavailability activates LOXL2 in the vascular ECM^37^, and inflammation leads to LOXL2 activation in fibrosis^45, 62, 63^. As endothelial dysfunction/loss of NO bioavailability and inflammation are key features of Ang II induced hypertension^64, 65^, these represent plausible mechanisms of LOXL2 activation in the hypertensive aorta and remain to be tested. The putative role of NO production (phospho eNOS/eNOS protein expression) and LOXL2 regulation by NO remain to be studied. This will be particularly important in females as the connection of LOXL2 to vasoconstriction/relaxation pathways and endothelial function were noted to be affected.

Interestingly, LOXL2’s catalytic function was found to be an important determinant of VSMC orientation in response to cyclic loading. This may be due to full-length LOXL2’s ability to deposit collagen IV, which is known to promote cell orientation/ polarity^66^ and promote VSMC differentiation. The elevated processing of LOXL2 and resultant accumulation of collagen I would promote VSMC hyperplasia and medial disorganization as noted in hypertensive WTs.

Our study has several limitations. In female mice, it is possible that PWV elevation arises with a sustained duration of hypertension that was not captured in the 3-week timeframe of our study. While we examined VSMCs and ECs in this study, adventitial fibroblasts are involved in adventitial remodeling in hypertension. Thus, fibroblasts as another source of LOXL2 in the vasculature and whether these respond to biomechanical strain also remains to be tested. The HAEC and HASMC used in this study were from males, and any cell autonomous effects of female sex were not probed and are a limitation of this study. The role of NO and inflammation in LOXL2 regulation and matrix remodeling in hypertensive males and females remain to be examined. Finally, as both aging and menopause are known to accelerate arterial stiffening independent of hypertension, the combined impact of aging/menopause and hypertension on male and female aortic mechanics is an important aspect of aging and female health and remain to be tested.

In conclusion, when considering the timeline of hypertension development, in vivo stiffness, passive aortic mechanics, and vascular reactivity results together, significant differences emerged between the sexes and illustrate that 1) ECM remodeling is a key feature of hypertension induced aortic stiffening in males and 2) the preservation of PWV in hypertensive females is due to the complex interplay between active and passive properties of the aorta. Our study further shows an important role for LOXL2 in the hypertensive aorta. It is well-established that hypertension elicits a reactive remodeling of the aorta in order to maintain matrix integrity as the aorta faces increasing pressure, but that sustained remodeling is detrimental to vascular health. Our study shows that LOXL2 depletion improves vascular load bearing in both males and females. Further, LOXL2 depletion confers clear benefits in male hypertensives and net benefits in females. Thus, LOXL2 represents a novel target to maintain aortic stiffness and aortic loadbearing in hypertensives that can circumvent the issues noted with inhibition of prototypic LOX, and this warrants further investigation.

## Supplemental Figures

**Supplemental Figure S1.** Systolic BP, Diastolic BP and mean arterial pressure (MAP) for all groups. n = 8-10 for each group.

**Supplemental Figure S2.** PWV and heart rate during PWV measurement for all groups. n = 8-10 for each group.

