## Supplementary figures and images for "Sex Differences and Role of Lysyl Oxidase Like 2 (LOXL2) in Angiotensin II-Induced Hypertension in Mice"

### Supplemental Figures 1-2

#### Supplemental Figure S1

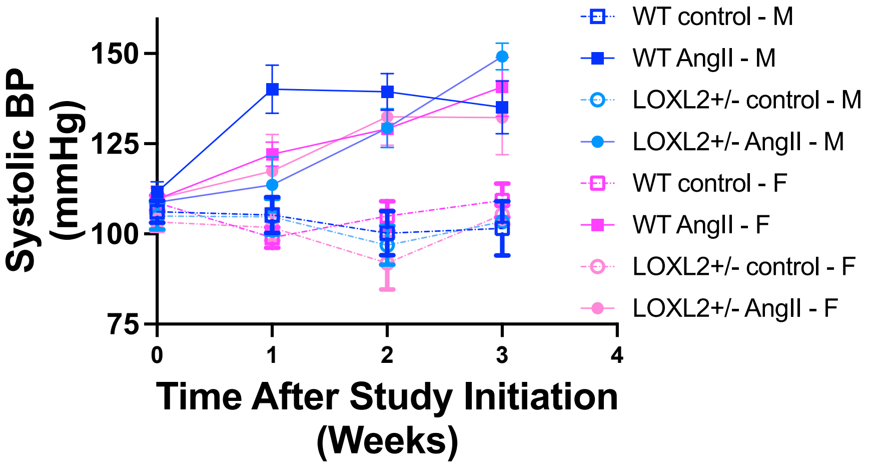


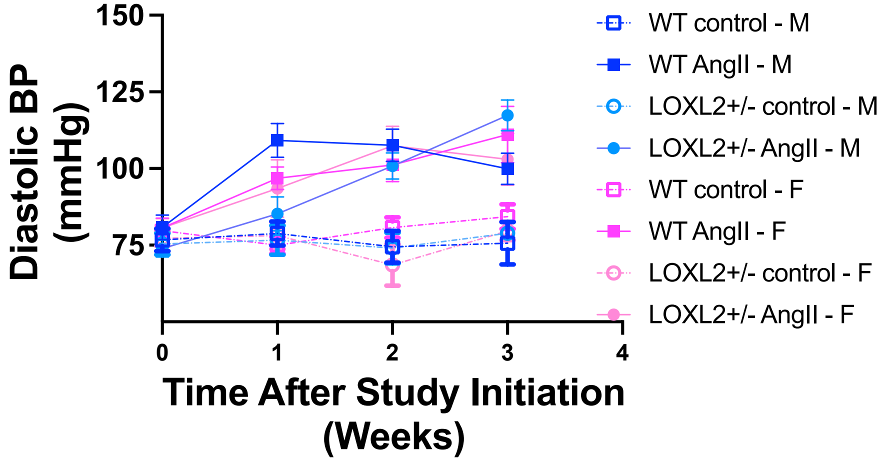


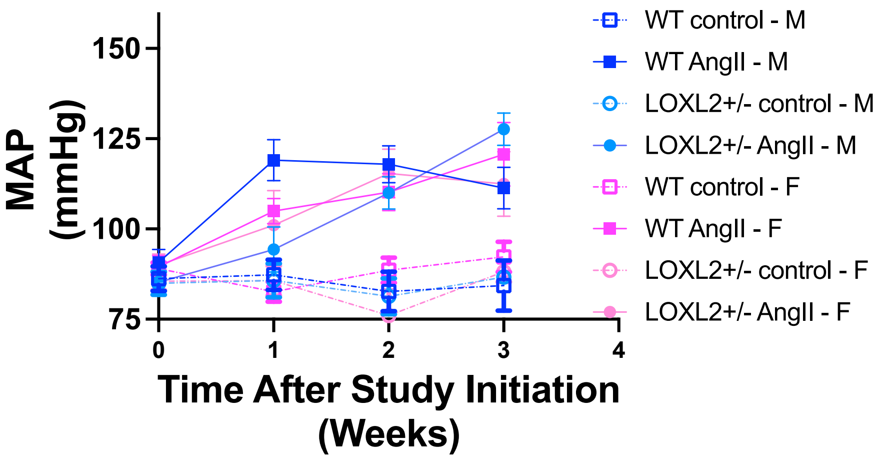

#### Supplemental Figure S2

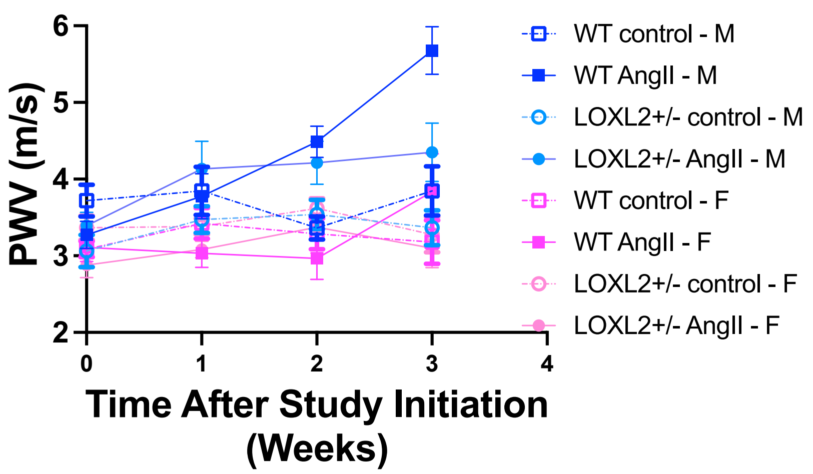


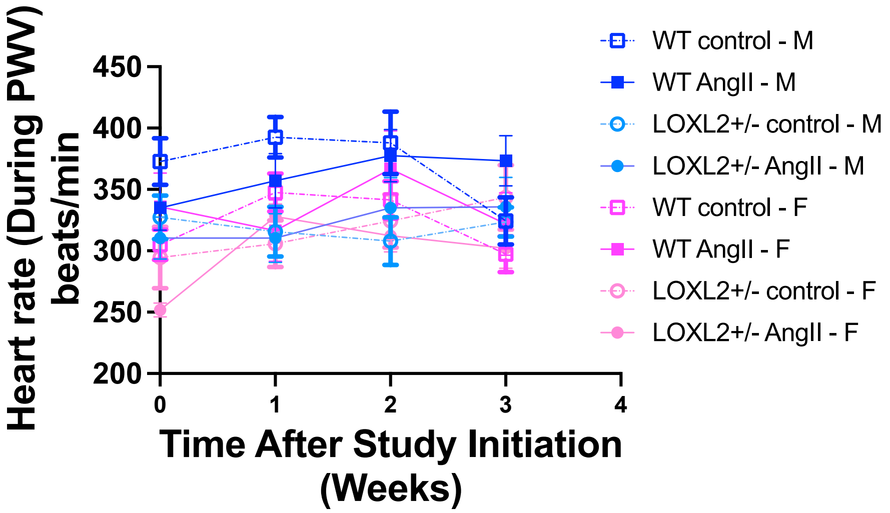
